# Deciphering Spatial Domains by Integrating Histopathological Image and Tran-scriptomics via Contrastive Learning

**DOI:** 10.1101/2022.09.30.510297

**Authors:** Yuansong Zeng, Rui Yin, Mai Luo, Jianing Chen, Zixiang Pan, Yutong Lu, Weijiang Yu, Yuedong Yang

## Abstract

Recent advances in spatial transcriptomics have enabled measurements of gene expression at cell/spot resolution meanwhile retaining both the spatial information and the histopathological images of the tissues. Deciphering the spatial domains of spots in the tissues is a vital step for various downstream tasks in spatial transcriptomics analysis. Existing methods have been developed for this purpose by combining gene expression and histopathological images to conquer noises in gene expression. However, current methods only use the histopathological images to construct spot relations without updating in the training stage, or simply concatenate the information from gene expression and images into one feature vector. Here, we propose a novel method ConGI to accurately decipher spatial domains by integrating gene expression and histopathological images, where the gene expression is adapted to image information through contrastive learning. We introduce three contrastive loss functions within and between modalities to learn the common semantic representations across all modalities while avoiding their meaningless modality-private noise information. The learned representations are then used for deciphering spatial domains through a clustering method. By comprehensive tests on tumor and normal spatial transcriptomics datasets, ConGI was shown to outperform existing methods in terms of spatial domain identification. More importantly, the learned representations from our model have also been used efficiently for various downstream tasks, including trajectory inference, clustering, and visualization.

## 1. Introduction

Recent advances in spatial transcriptomics (ST) technologies have enabled measurements of gene expression at cell/spot resolution meanwhile retaining both the spatial coordinates and the histopathological image of the tissue, such as Visium from 10X Genomics, XYZeq [1], and Slide-seq [2]. Most existing ST technologies could sequence gene expressions at each spot that contains two to dozens of cells [3]. These effective ST technologies have transfigured our perspectives on the developing amyotrophic lateral sclerosis, human heart, Alzheimer’s disease, and the human squamous cell carcinoma [4–7]. Deciphering spatial domains is one of the great challenges of spatial transcriptomics analysis. For example, cells often residing within different cortical layers of the human cerebral cortex differ in morphology, gene expressions, and physiology, and the laminar organization of the human cerebral cortex is related to its biological functions [8]. In fact, spatial domain identification could be resolved by applying clustering methods [9–12]. Nevertheless, conventional clustering methods are meeting grand challenges since they do not incorporate the unique properties of spatial transcriptomics or they are difficult to deal with the sparsity and high dimensional features of gene expression data [13, 14].

To resolve these challenges, a wide variety of clustering algorithms have been developed for ST analysis [15–19]. Several early spatial clustering methods account for the spatial dependency of gene expression by considering the similarity between adjacent spots. For example, BayesSpace [20] is a Bayesian statistical model that promotes the nearest spots to belong to the same cluster via introducing spatial neighbor structure into the prior. Similarly, Giotto [21] deciphers spatial domains by applying a hidden Markov random field (HMRF) model that integrates the spatial neighbor information. However, they achieve limited performance since they don’t capture the non-linear characteristics of gene expression. Therefore, a few deep learning-based methods have been designed for learning non-linear relationships. For example, SEDR [22] embeds the spatial structure of ST via a variational graph autoencoder network and simultaneously learns latent gene representations through the autoencoder network. Although these aforementioned methods consider the spatial structure of ST, they pre-defined the similar relations between spots without updating in the late stage. Therefore, STAGATE [23] is proposed for adaptively integrating the gene expression profiles and structure of ST via an attention graph convolutional network [24]. Even though these structure-based methods achieve decent performance on some datasets by depending on the spatial structure and gene expression, their performance may be influenced when the gene expressions contain a mass of noises due to the defect of the current sequencing techniques [25, 26]. Previous methods show that histopathological images can be used for predicting gene expression[13, 27], indicating images also contain abundant information. Thus, it is promising to conquer noises in gene expression by introducing image features as complementary information for gene expression profiles.

To take full use of the information from images, several methods have been developed. For example, stLearn [28] computes the morphological distance between spots by using their corresponding features extracted from the histopathological image and applies these distances and the spatial structure to smooth gene expression. SpaGCN [29] uses both the histopathological image and spatial coordinates to build relations between spots, which are then fed into a graph convolutional layer [30] to propagate gene expression information between neighboring spots. conST [31] first concatenates gene expressions and the pre-extracted morphology features that are extracted from images via the Masked Auto-encoder (MAE) into a feature vector [32], which is then fed into a graph convolutional network to learn latent representations.

Though these methods have been designed for integrating the gene expression and histopathological images, they use the histopathological images to construct spot relations without updating in the training stage, or simply concatenate the information from image and gene expression into one feature vector. These would cause suboptimal performance since they may learn meaningless modality-private noise information of each modality, introducing additional noises to domain clustering. Yet, this can be avoided by applying cross-modal contrastive learning, which is able to learn the common semantics across all modalities while reducing their meaningless modality-private noise information [33].

To this end, here, we propose a novel method ConGI to accurately decipher spatial domains by integrating gene expression and histopathological images, where the gene expression is adapted to image information through contrastive learning. The natural rationale of our method is to leverage the correspondence between gene expression and cellular phenotypical information at the spot level. To learn the common semantic representations across all modalities while avoiding their meaningless modality-private noise information, we introduce three contrastive loss functions within and between modalities, including gene expression to gene expression, image to image, and image to gene expression. Next, the learned representations are then used for deciphering the spatial domains through clustering methods. By comprehensive tests on tumor and normal datasets, ConGI was shown to outperform existing methods in terms of spatial domain identification. More importantly, the learned representations from our model have also been used efficiently for various downstream tasks, including trajectory inference, clustering, and visualization.

## 2. Materials and Methods

### 2.1 Datasets and pre-processing

To evaluate the performance of our method, we employed four spatial transcriptomics datasets, including the human HER2-positive breast tumor dataset (HER2+) [34], the human dorsolateral prefrontal cortex dataset (spatialLIBD) [8], the human epidermal growth factor receptor (HER) 2-amplified (HER+) invasive ductal carcinoma (IDC) sample (https://sup-port.10xgenomics.com/spatial-gene-expression/datasets), and the mouse brain datasets (https://www.10xgenomics.com/resources/datasets). The HER2+ dataset was measured by spatial transcriptomics technology (ST technology), which included eight tissue sections with annotations from pathologists. We removed section C in the HER2+ due to the number of spots being less than 200. The spatialLIBD consisted of 12 sections that were measured by the 10x Visium platform. The mouse brain dataset contained one anterior section generated by the 10x Visium platform. The IDC dataset included one section generated by the 10x Visium platform. Each dataset contained histopathological images, gene expression at the spatial spots, and their corresponding coordinates. For each histopathological image, we cropped (W × H) pixels around each spot, where H and W were the height and width of image patches, respectively. Both W and H were set to 112, matching the diameter of each spot. We provided two strategies to pre-process the gene expression data of each tissue section. For the dataset generated by 10x Visium, we used PCA to reduce the dimension of gene expression to 300 as recommended in ref [29, 31]. For other datasets, we followed ref [13] to select the top 1000 highly variable genes; For a given spot, its counts were divided by the total counts for the given spot and multiplied by the scale factor of 1,000,000. This was then natural-log transformed via log (1+x), where *x* was the normalized count.

### 2.2 The Architecture of ConGI

ConGI is a deep learning-based method for deciphering spatial domains by integrating histopathological images and spatial transcriptomics via contrastive learning. To achieve this, as illustrated in Figure 1, given pairs of spatial transcriptomics (gene expression) and the image patch cropped from histopathological images at each spot, we apply two independent encoders (a CNN f^i^ and an MLP f^*g*^) to learn the low-dimensional representations via contrastive learning. Concretely, we first distort the paired image patch and gene expression data slightly by adding noises to them through their corresponding data augmentation techniques. The augmented data sets are then fed into the encoders for images and gene expressions to learn the low-dimensional representations, separately. These learned representations are then projected into the space where contrastive loss is applied to jointly learn from pairwise data via three contrastive learning losses, including gene expression to gene expression (*L*_g2g_), image to image (*L*_i2i_), and image to gene expression (*L*_i2g_). ConGI pulls the pairwise data within and between modalities together and contrasts the unmatching pairs apart. After training, the low-dimensional representations of gene expression and image patch were combined together for clustering via the mclust package. More importantly, the learned representations have been used efficiently for various downstream tasks, including trajectory inference, clustering, and visualization.

**Figure 1.**
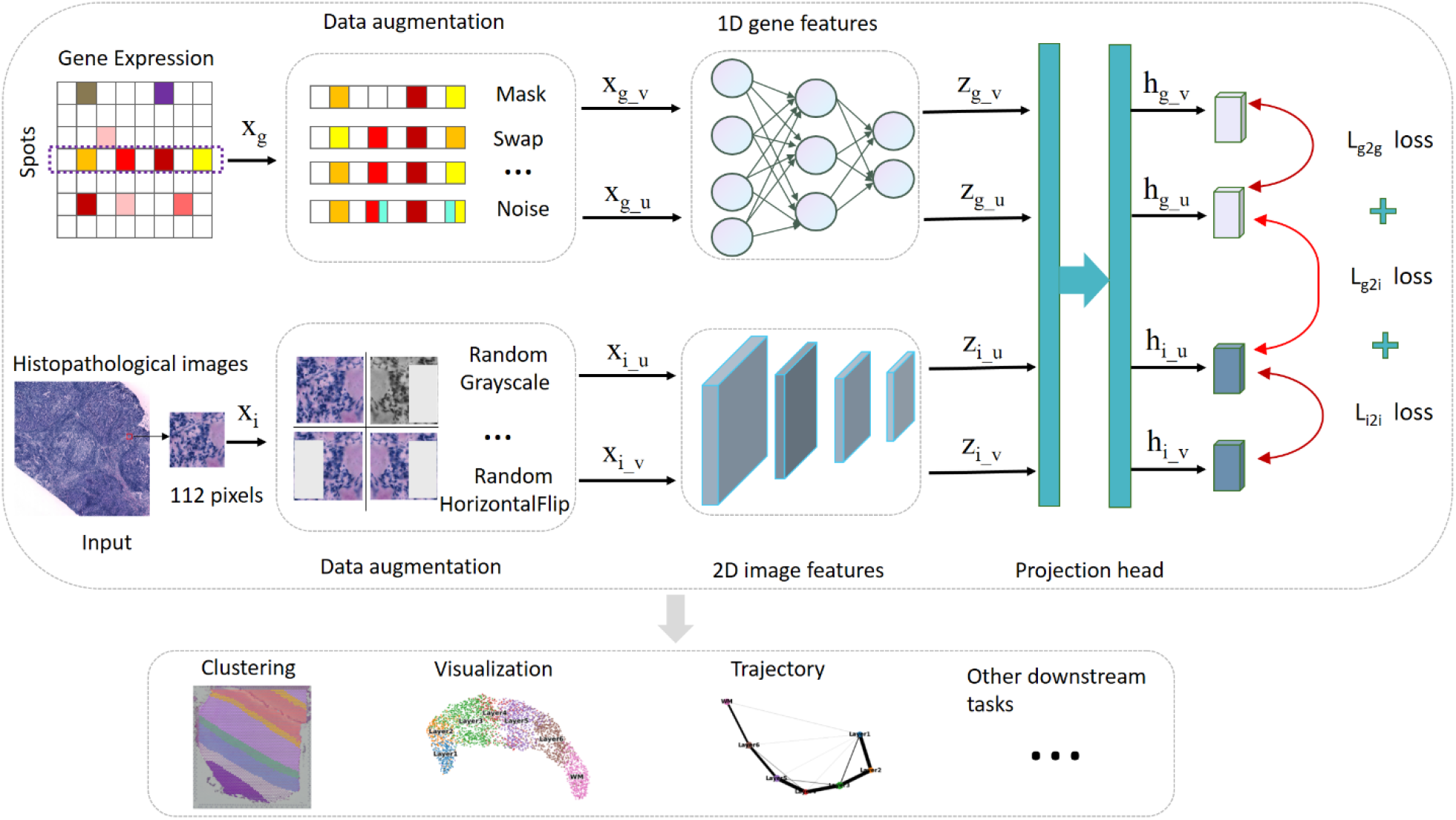
The schematic overview of the ConGI for deciphering spatial domains by integrating image and gene expression. The input of ConGI is the paired image patch and gene expression of a spot. We first distort the paired image patch and gene expression data slightly by adding noises to them through their corresponding data augmentation techniques. The augmented data sets are then fed into encoders for image and gene expression to learn the low-dimensional representations, respectively. These learned representations are then projected into the space where contrastive loss is applied to jointly learn from pairwise data via three contrastive learning losses, including gene expression to gene expression (*L*_g2g_), image to image (L_i2i_), and gene expression to image (L_*g*2*i*_). Finally, the learned representations are then used for deciphering spatial domains, visualization, and trajectory inference.

#### 2.2.1 Encoders for gene expression and image

##### Gene expression encoder

The gene expression encoder is used for extracting the low-dimensional representations of gene expression X ∈ ℝ^*n*×*d*^, where *n* and *d* are the numbers of spots and the feature dimension of the spot. The gene expression encoder f^*g*^ is a neural network-based model applying a multi-layer perceptron (MLP) with fully-connected layers and non-linearities. Specifically, for a given spot, we first generate two augmented gene expressions x_*g_u*_ and x_*g_v*_ by distorting the original gene expression x_g_ ∈ X of the spot through data augmentation techniques, such as the random mask and random swap. The augmented data sets are then fed into gene expression encoder f^*g*^ to obtain the corresponding latent vectors z_*g_u*_ and z_*g_v*_ as follows:

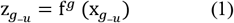

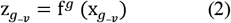

##### Image encoder

The image encoder f^*i*^ aims to capture the morphological feature of each spot from the image patch. Here, the backbone of the image encoder f^*i*^ is a classic convolutional neural network (CNN) DenseNet121 with the pre-trained ImageNet weights. In our setting, we reserve the pretrained weights of DenseNet121 due to the limited training images. The input of the image encoder is the image patch cropped from the whole histopathological image, which is only matching to the corresponding gene expression. Similarly, we also generate two augmented image patches x*_i_-u__* and x_*i*_-*v*__ by distorting the original image patch x_i_ through data augmentation techniques, such as RandomGrayscale and Random HorizontalFlip. The augmented data sets are then fed into the image encoder f^*i*^ to obtain the corresponding latent vectors z_*i*_-*u*__ and z_*i*_-*v*__ as follows:

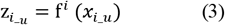

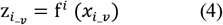

##### Projection head

For learning the common semantic information from image and gene expression, we apply a small neural network projection head pg (.) layer to take the gene expression and the image representations into a shared space. Here, the pg (.) layer consists of a two-layei neural network, which connect directly with the image and gene expression encoders. The pg (.) projects the latent features z_*g-u*_ and z_*g-v*_ ofgene expression and the latent features z_i_-u__ and z_<u>i</u>_-*u*__ of the image patch into a shared space as follows:

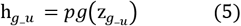

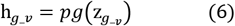

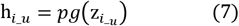

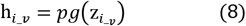

#### 2.2.2 Contrastive learning loss functions

To learn the common semantic representations between images and gene expressions while avoiding their meaningless modality-private noise information, we introduce three contrastive learning losses within and between modalities. The contrastive learning between modalities aims to pull paired low-dimensional representations h_*i_u*_ and *h_g_u_* of image patch and gene expression together while contrasting those unmatching pairs apart. To better learn the characteristics of each modality, we conduct contrastive learning within the modality for image and gene expression, respectively. They pull their pairwise data together while pushing those unmatching pairs apart. For achieving these goals, we design the contrastive learning loss function following SimCLR [35], which is a simple framework for contrastive learning of visual representations. SimCLR achieves SOTA performance on many datasets. Concretely, we randomly build a minibatch of N spots and define the contrastive prediction task on paired spots derived from the minibatch, resulting in 2N paired spots. We do not build negative spots explicitly. By contrast, similar to ref [36], given a positive pair, we take the other 2(N-1) spots within a minibatch as negative spots. Thus, the within and between contrastive loss functions for positive pairs 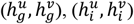, and 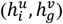 can be defined as follows:

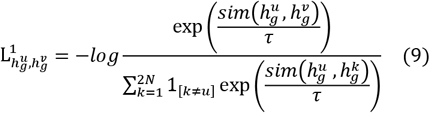

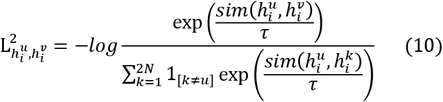

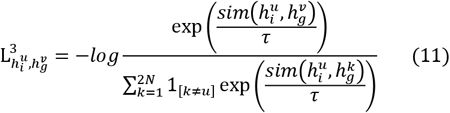

where 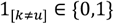 is an indicator function evaluating to 1 if k ≠ *u*, and *τ* means the temperature parameter. The term 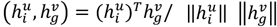 represents the dot product between 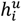 and 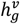. Each contrastive loss is calculated across all positive pairs, i.e., both 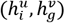 and 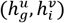 in a mini-batch. Thus, three contrastive losses can be formulated as follows:

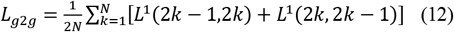

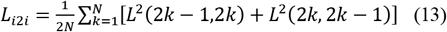

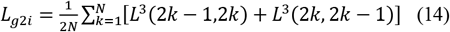

Finally, the total loss of our model can be summed as follows:

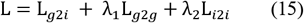

where λ_1_ *and* λ_2_ are hyper-parameters used for controlling the contribution of losses L_*g*2*g*_ *and L*_*i*2*i*_ for the final loss. For all datasets, λ_1_ = λ_2_ = 0.1.

After the model is trained, the final representation of each spot can be defined as follows:

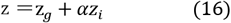

where *α* is a hyper-parameter used for controlling the contribution for the final representation of each spot. For all datasets, *α* = 0.1.

#### 2.2.3 Clustering

We take different strategies to identify spatial domains using learned representations from ConGI. When the number of spatial domains is specific, we follow ref [23] to apply the mclust [37] clustering algorithm to decipher spatial domains. For datasets without the number of clusters, we identify spatial domains through the Louvain algorithm implemented in the popular package scanpy. After clustering, we follow SpaGCN to provide an optional refinement step for the clustering results. In this step, for a given spot, we reassign its label to the same spatial domain as the primary label of its neighboring spots if more than half of its neighboring spots are assigned to a different domain. We perform cluster refinement for all datasets.

### 2.3 Hyper-parameters setting

The ConGI was implemented in python and PyTorch. For the gene expression encoder, the dimensions of hidden layers were set to [128,128]. The image encoder used the DenseNet121 with default pre-trained weights from torchvision.models. The dimensions of the projection head were set to [128,128]. Our models were optimized via the AdamW optimizer with a learning rate of 0.003. The training batch size was set to 32 when the total number of spots was less than 1000. In other situations, the training batch size was set to 64. In the processing of the refinement step, the number of neighboring spots was set to 24 when the number of total spots was greater than 1000. Otherwise, the number of neighboring spots was set to 4. All results reported in this paper were conducted on Ubuntu 18.04.7 LTS with Intel^®^ Core (TM) i7-8700K CPU @ 3.70 GHz and 256 GB memory.

### 2.4 Evaluation criteria

#### 2.4.1 Clustering performance

Three common clustering metrics are applied for evaluating clustering results in this study including Normalized Mutual Information (NMI) [38], Adjusted Rand Index (ARI) [39], and Clustering Accuracy (CA)[40].

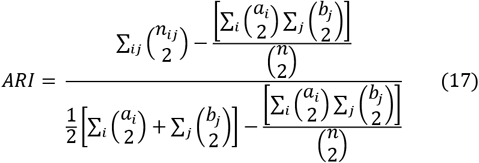

where a_*i*_ and b_*j*_ are the number of samples appearing in the *i-th* predicted cluster and the *j-th* true cluster, respectively. n_*ij*_-means the number of overlaps between the *i-th* predicted cluster and the *j-th* true cluster.

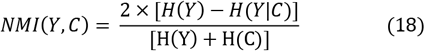

where C and Y are the predicted clusters and the real clusters, respectively. The function H () is used for calculating the entropy.

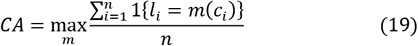

where *n* is the entire number of samples, and *m* ranges over all probable one-to-one mapping between clustering assignment *c_i_*, and true label *l_i_*.

### 2.5 Benchmark methods

To evaluate the performance of our method, we compared ConGI with other tools including STAGATE, scanpy, conST, SpaGCN, SEDR, stLearn, BayesSpace, Giotto, and Seurat [41]. For all competing methods, we used the default hyper-parameters and pre-processing for datasets recommended in the original paper to test all datasets. For methods scanpy and Seurat, we used the default value of parameter “resolution” to determine the number of clusters. For other methods, we fed them with the number of true clusters for clustering.

## 3. Results

### 3.1 Application to spatially resolved tumor sample transcriptomics

To demonstrate the performance of our method on the spatially resolved tumor sample transcriptomics, we analyzed the human HER2-positive breast tumor dataset (HER2+) and the human epidermal growth factor receptor (HER)2-amplified (HER+) invasive ductal carcinoma (IDC). We first compared the performance of our method with competing methods on the HER2+ data. As shown in Figure 2(a), ConGI outperformed all competing methods in terms of the average ARI. Specifically, the average ARI of our method was 11.2% higher than the second-ranked method Giotto. stLearn obtained a decent performance with an average ARI value of 0.22. SpaGCN and conST achieved similar performance. BayesSpace and STAGATE achieved similar clustering results, both of them performed better than SEDR. Interestingly, methods (i.e., scanpy, SEDR, and STAGATE) only using the gene expression information performed worse than methods (i.e., ConGI, stLearn, conST, and SpaGCN) considering image information. These results demonstrated that image features were beneficial for clustering spatial domains. To further confirm the superior results, we compared the manually annotated regions for section E1. Figure 2(b) and Supplementary Figure S1 showed that our method revealed spatial domains that agreed better with the manually annotated tissue regions than competing methods. Although stLearn and conST also utilized histopathological images, they still assigned the wrong spatial domains for most spots. When the HER2+ dataset was tested by the other metrics (NMI and CA), a similar trend could be found (Supplementary Figure S2).

**Figure 2.**
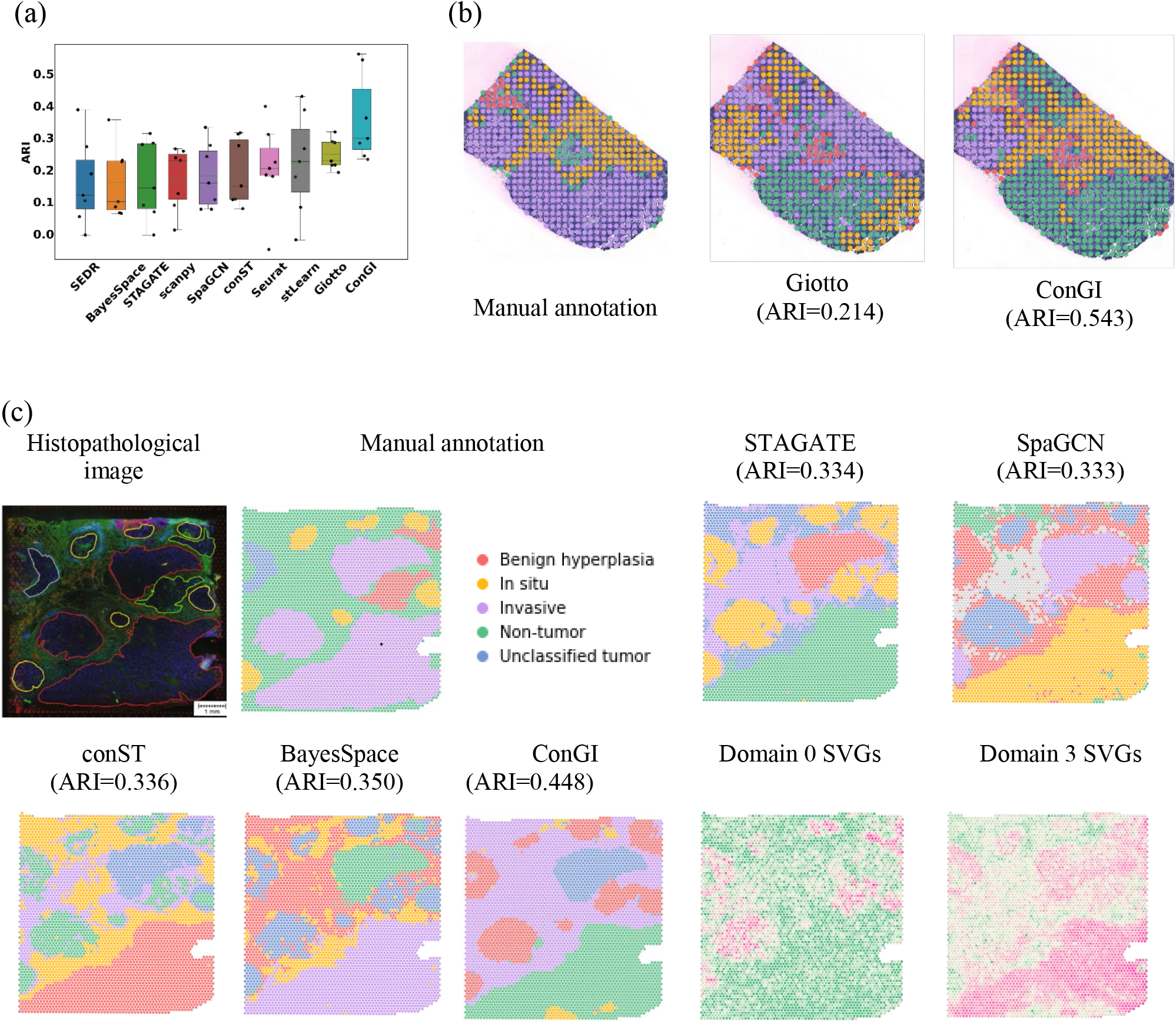
Spatial domains and SVGs detected in the human HER2-positive breast tumor (HER2+) dataset and the invasive ductal carcinoma (IDC) dataset. (a) Boxplot of clustering accuracy in all sections of the HER2+ dataset in terms of ARI values for all methods. (b) Manually annotated regions of section E1 of the HER2+ dataset, spatial domains detected by Giotto and ConGI. (c) The histopathological image and manually annotated regions for the IDC data, spatial domains detected by STAGATE, SpaGCN, conST, BayesSpace, and ConGI, and the spatial expression patterns of SVGs for ConGI predicted spatial domains 0 (FAM234B) and 3 (MUC1).

We further evaluated the performance of our model on the IDC dataset with the nearly single cell super-resolution. Pathologists identified regions of benign hyperplasia, predominantly invasive carcinoma (IC), and carcinoma in situ, which were used as ground truth labels to evaluate the clustering accuracy. As shown in Figure 2(c) and Supplementary Figure S3, our model achieved the best clustering accuracy with an ARI of 0.448, which was 9.8% higher than the second method BayesSpace. Methods STAGATE, SpaGCN, conST, and Seurat achieved similar results with an ARI value of around 0.330. Both SEDR and stLearn achieved similar performance with an ARI value of about 0.286. scanpy achieved the lowest performance with the ARI value of 0.231. Compared to the ground truth labels, our method could identify all tumor regions and non-tumor regions. Though ConGI failed to identify in situ and benign hyperplasia regions, other competing methods also mixed them together. This is maybe because it was difficult to distinguish regions between in situ and benign hyperplasia. To further confirm that ConGI could investigate the biological relevance, we detected the spatially variable genes (SVGs) for the IDC dataset via the same detection strategies used in SpaGCN. The results showed that FAM234B enriched in domain 0. Similarly, MUC1 was enriched in domain 3 in the histopathological image. The results demonstrated that ConGI showed similar biological tissue patterns to manual annotations.

### 3.2 Application to spatially resolved normal sample tran-scriptomics

To evaluate the performance of our method on the spatially resolved normal sample transcriptomics, we analyzed a human dorsolateral prefrontal cortex (spatialLIBD) dataset and a mouse brain anterior tissue. We first analyzed the spatialLIBD consisting of 12 tissue slices from the human dorsolateral prefrontal cortex in three human brains, which spanned six neuronal layers and white matter. As shown in Figure 3(b), our method outperformed other methods in terms of the average ARI. Concretely, the average ARI of our method was 4.82% higher than the second-ranked method STAGATE. Though SEDR achieved decent performance and ranked third, the average ARI value was much lower (13.42%) than that of our method. The performance of other image-based methods (such as SpaGCN and stLearn) didn’t outperform SEDR, which may be because they couldn’t deal with noises generated from the image patch. These results demonstrated that our method could efficiently learn the common semantics across all modalities while avoiding their meaningless modality-private noise information. BayesSpace and Seurat achieved similar performances and ranked seventh and eighth, respectively. Giotto obtained the lowest performance. To further confirm the better results, we compared the manually annotated layer structure for section 151509, which had clear layer boundaries. Figure 3(c) and Supplementary Figure S4 showed that our method revealed spatial domains that better agreed with the manually annotated tissue layers (Figure 3a) than competing methods. Although the results of STAGATE had seen seven relatively clear layers, most spots in Layer2 mixed with spots in Layer1 together. Most spots were mixed wrongly in the spatial domains predicted by scanpy and Giotto. In addition, we listed the results of all sections measured by NMI and CA in Supplementary Figure S5, a similar trend could be found. We further identify the spatially variable genes (SVGs) for section 151509. As shown in Figure 3(c), HPCAL1 was enriched in domain 4. Similarly, MBP was enriched in domain 7 in the histopathological image. The results demonstrated that ConGI showed similar biological tissue patterns to manual annotations.

**Figure 3.**
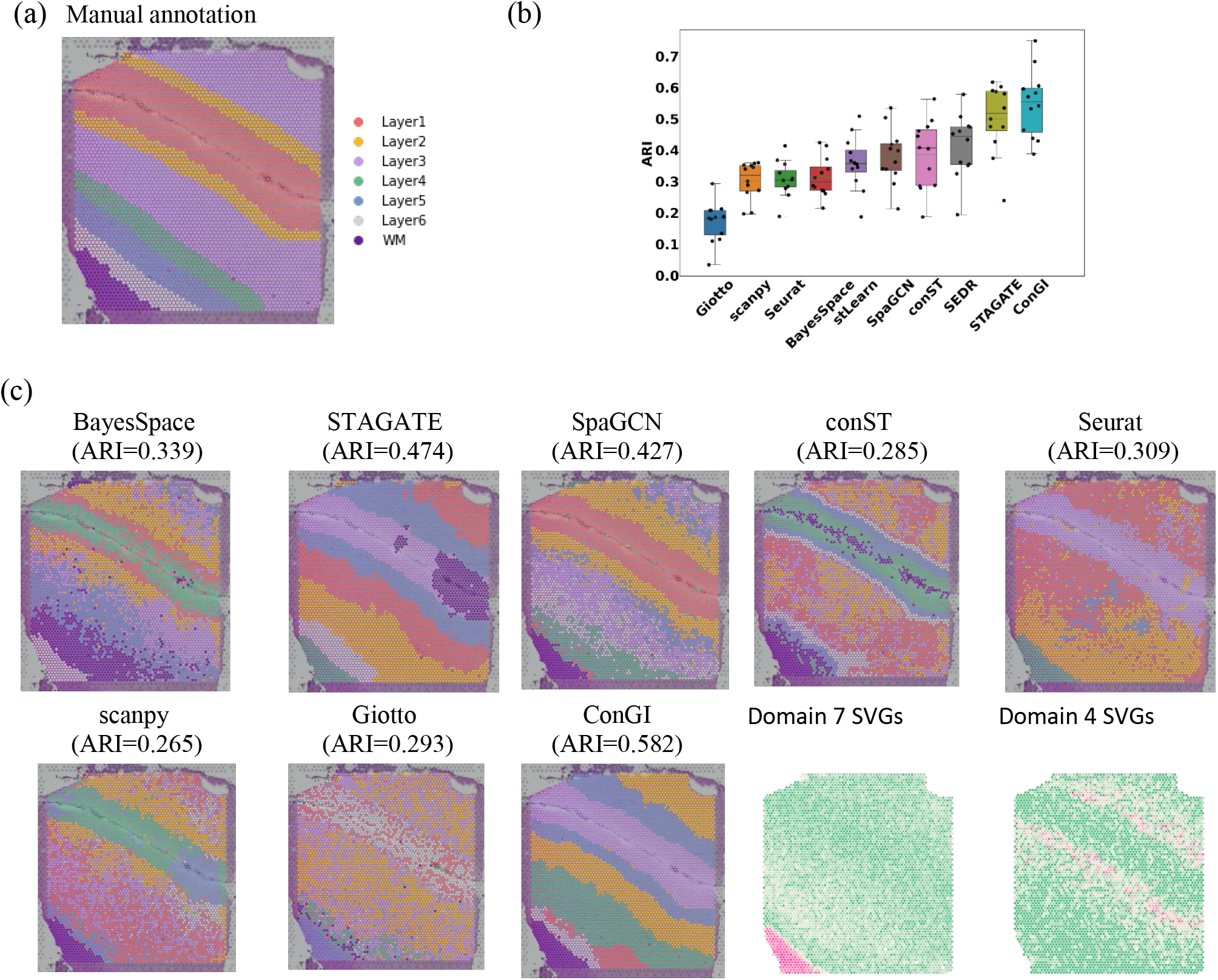
Spatial domains and SVGs detected in the human dorsolateral prefrontal cortex dataset (spatialLIBD). (a) Histopathological image of the tissue section 151509 with manually annotated layer structure. (b) Boxplot of clustering accuracy in all sections of the spatialLIBD dataset in terms of ARI values for all methods. (c) Spatial domains were detected by BayesSpace, STAGATE, SpaGCN, conST, Seurat, scanpy, Giotto, and ConGI in section 151509, respectively, and the spatial expression patterns of SVGs for ConGI predicted domain 4 (HPCAL1) and domain 7 (MBP).

We further analyzed the mouse brain anterior tissue, which had more complex tissue structures than the spatialLIBD dataset. The manual annotations from pathologists were used as the ground truth for evaluating the clustering performance. As shown in Figure 4 and Supplementary Figure S6. Our method achieved the highest ARI value of 0.401. We found that the performance of our method was slightly higher than the second-ranked method Giotto. This is likely because the information from the image included less useful information for deciphering the domains. Actually, we did not find obvious boundaries among spatial domains in the histopathological image of the mouse brain anterior tissue. The other image-based method conST wrongly predicted all spatial domains into one domain. Seurat, SpaGCN, and STAGATE achieved similar decent results. However, some comparison methods achieved low clustering performance, such as SEDR obtained the ARI value of 0.28. The results showed that our method still could achieve a decent performance when there were not any clear boundaries between spatial domains in the histopathological image.

**Figure 4.**
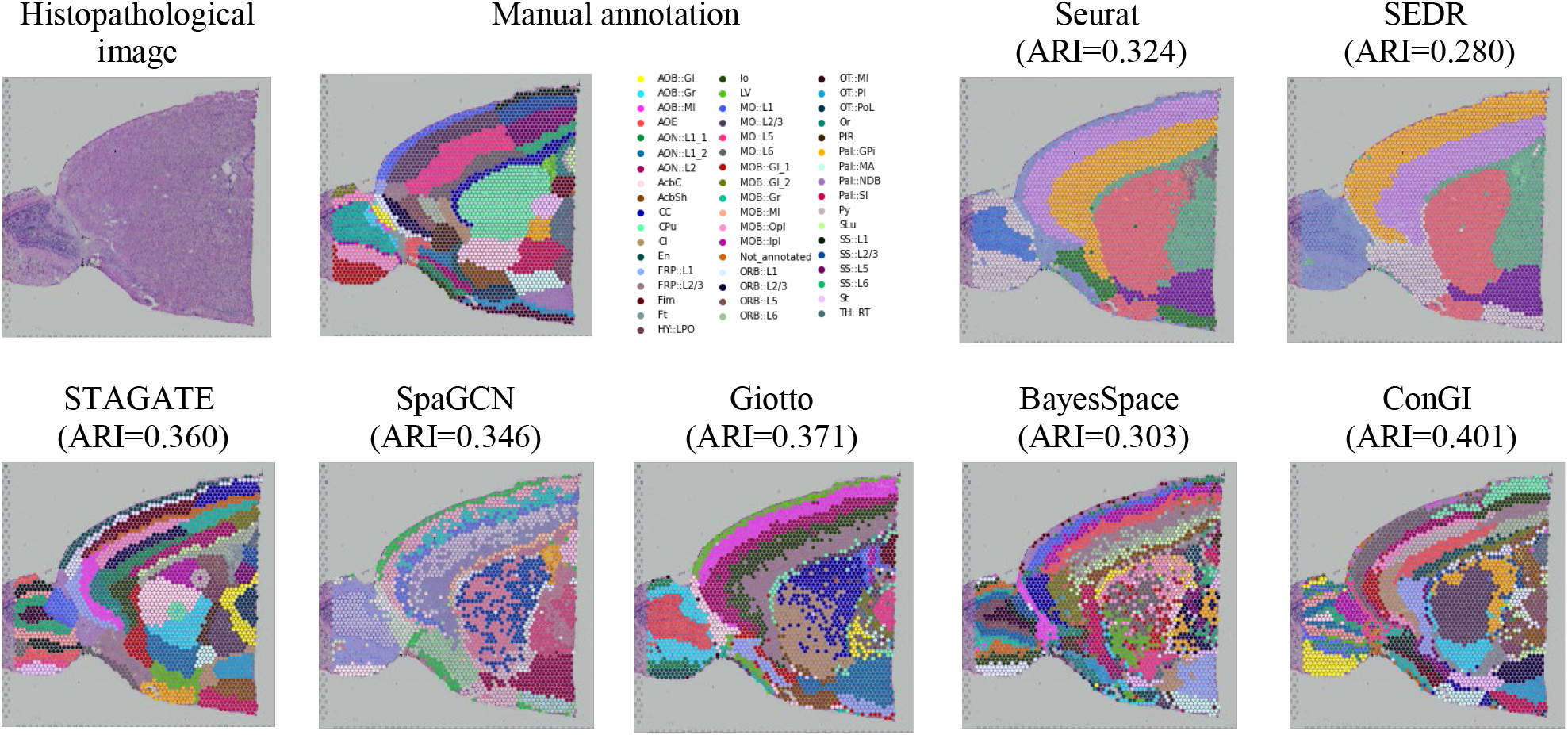
The histopathological image and manually annotated regions for the mouse brain anterior sample, spatial domains detected by Seurat, SEDR, STAGATE, SpaGCN, Giotto, BayesSpace, and ConGI.

### 3.3 ConGI learns effective latent representations from image and gene expression

To show the effective representations learned by our model could reveal the distance between spatial domains and depict the trajectory, we visualized the low-dimensional representations through the UMAP [42] in the 2D space and plotted the trajectory inference via the PAGA [43] algorithm employed in the package scanpy. We took tissue section E1 from the HER2+ dataset and tissue section 151509 from the spatialLIBD dataset as examples. We first evaluated our method in section E1. As shown in Figure 5(a) and Supplementary Figure S7, STAGATE and BayesSpace completely mixed spots in regions of connective tissue, invasive cancer, and immune infiltrate. A similar trend could be found in the UMAP of other competing methods, such as Giotto and SpaGCN. In contrast, our method separated explicitly the domains invasive cancer and immune infiltrate. Though ConGI didn’t separate the immune infiltrate and the connective tissue, other methods were also difficult to distance them. This is likely because they were difficult to distinguish. A similar trend could be found when visualizing the tissue section 151509 of the spatialLIBD dataset. As shown in Figure 5 (b) and Supplementary Figure S8, In the UMAP plots of BayesSpace, scanpy, and Seurat, most spots of different layers were mixed, such as spots from layers 2 to 4. In the UMAP plots of conST, spots from layer 1 and spots from layer 6 were mixed. However, our method and STAGATE could well separate most spots in different layers. Since section 151509 contained explicitly the layer structure of the human dorsolateral prefrontal cortex, we further validate the inferred trajectory based on the representations generated by all methods. The PAGA graphs of both ConGI and STAGATE representations showed a nearly linear development trajectory from layer 1 to layer 6 as well as the similarity between adjacent layers. Nevertheless, the PAGA results of BayesSpace showed that most layers were mixed. Layer 6 and layer 4 were mixed in the PAGA results of SpaGCN. The results of UMAP and trajectory inference demonstrated that our method could learn effective representations for downstream tasks.

**Figure 5.**
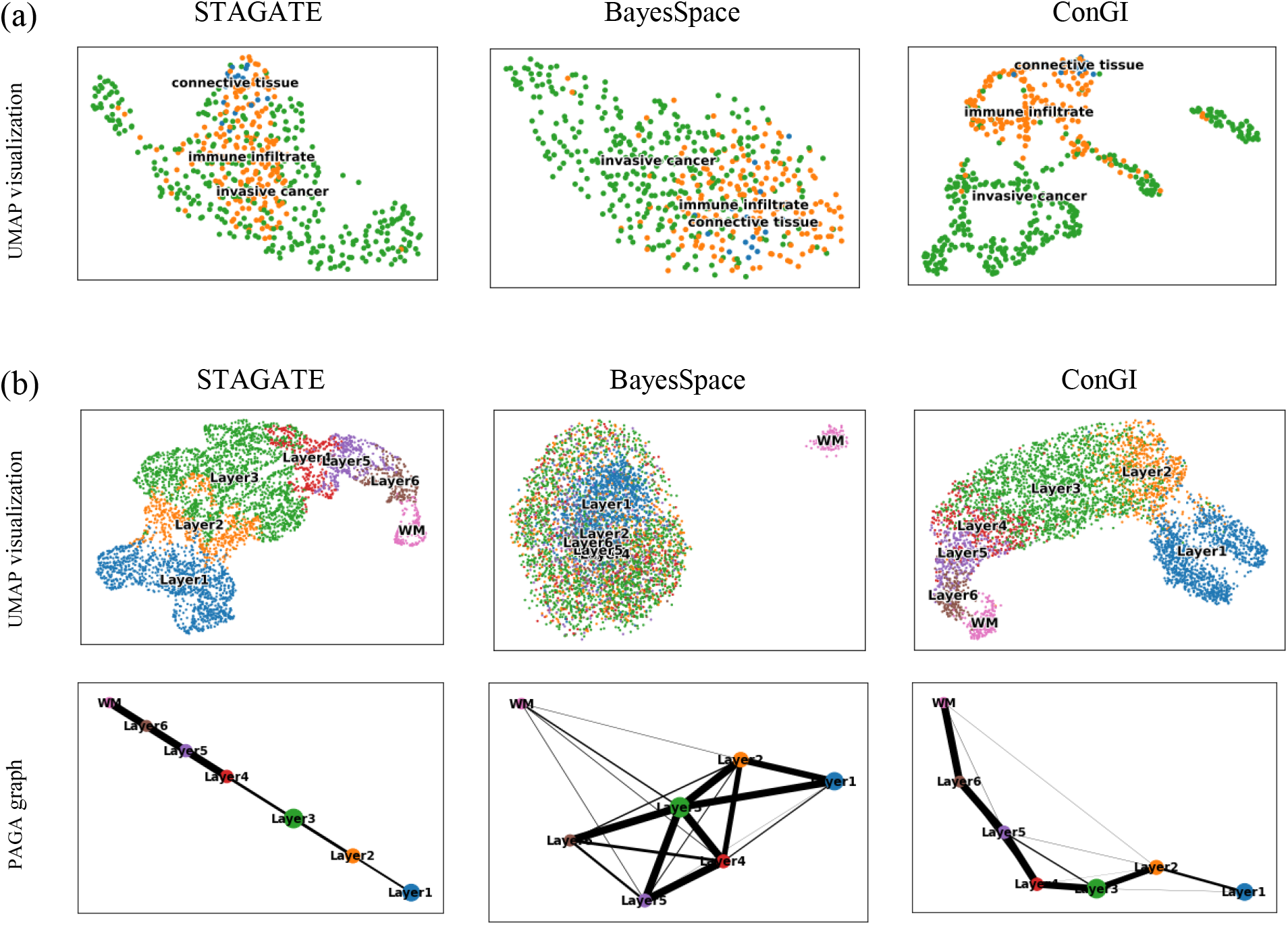
UMAP visualizations and PAGA graphs using representations generated by STAGATE, BayesSpace, and ConGI respectively (a) in the section E1 of the dataset HER2+ and (b) the section 151509 of the dataset spatialLIBD.

### 3.4 Ablation study

To investigate the contributions of components for our method ConGI, we conducted ablation studies on the HER2+ dataset. As shown in Figure 6, We first tested whether the performance of our model benefited by learning information adaptively from histopathological images. We found that the performance of our method decreased by 3% if we didn’t update the weights of the DenseNet121 through the histopathological images (domain knowledge) in the training stage. These changes demonstrated that the performance of our method was improved by learning information adaptively from histopathological images. The removal of the contrastive learning loss within gene expressions (L_g2g_) and images (L_i2i_) caused decreases of 1.5% and 2.4%, respectively. The removal of the contrastive learning loss between gene expression and the image (L_g2i_) caused a significant decrease of 11%, which indicated that our model was able to effectively learn the common semantics across image and gene expression and avoid their meaningless view-private noise information via contrastive learning. In conclusion, the better performance of our method relied on the cooperation of the modules.

**Figure 6.**
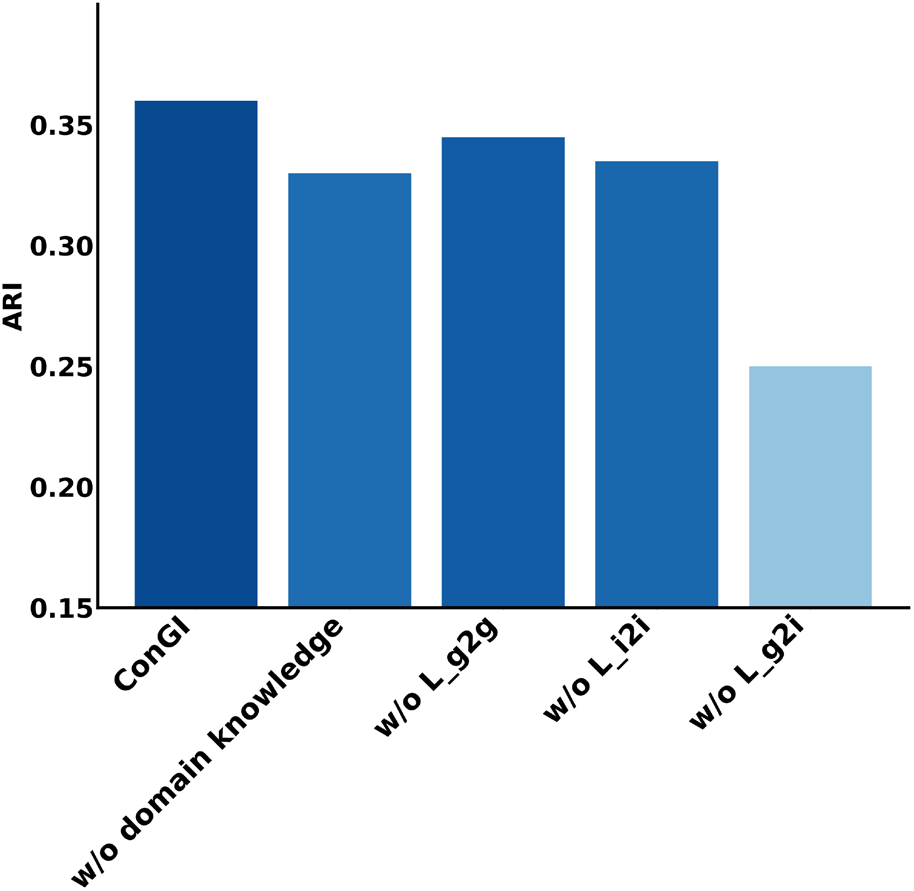
Ablation study for ConGI on the HER2+ dataset measured by ARI.

## 4. Discussion

Accurate identification of spatial domains is essential for researchers to understand tissue organization and biological functions. In this study, we proposed a novel method ConGI to accurately decipher spatial domains by integrating gene expression and the histopathological image, where the gene expression is adapted to image information through contrastive learning. The learned representations are then used for deciphering the spatial domains through a clustering method. By comprehensive tests on cancer and normal spatial transcriptomics datasets, ConGI was shown to outperform existing methods in terms of spatial domain identification. More importantly, the learned representations from our model have also been used efficiently for various downstream tasks, including trajectory inference, clustering, and visualization.

While a few methods, such as stLearn and SpaGCN have been developed for deciphering spatial domains by combining the gene expression and histopathological images to conquer noises in the gene expression. However, they only use the histopathological images to construct spot relations without updating in the training stage. This may lead to poor performance if the image features construct the wrong spot relationships. This is because the spot relations are not updated in the training phase. Though the other method conST also integrates directly the image features, it simply concatenates the information from gene expression and image into one feature vector as the input to train the model. This structure cannot accommodate the different processing needs of each modality and cannot avoid their meaningless modality-private noise information. In contrast, our model applies CNN-based and MLP-based models to learn low-dimensional representations from image and gene expression separately, where the gene expression is adapted to image information through contrastive learning. Especially, the image feature in our framework could be learned adaptively through contrastive loss, which is beneficial to avoid the noise information of images. ConGI can efficiently learn the common semantics across all modalities while avoiding their meaningless modality-private noise information. By comprehensive tests on cancer and normal datasets, ConGI was shown to outperform existing methods in terms of spatial domain identification.

In spite of the superior performance, ConGI can be improved in several aspects. Firstly, as a deep learning method, our model shows some limitations including the black-box nature of AI models, which can be addressed through downstream analysis such as spatial variable genes identification that can ameliorate some of the problems and bring insights into the cluster labels. Secondly, we use the DenseNet121 with pre-trained ImageNet weights for extracting the features from histopathological images while having not fully used the big models trained using the histopathological images in the field of biological medicine. With the relatively easy acquirement of histopathological images, it is promising to pre-train a specific big model on histopathological images and use it in our extraction. In conclusion, this study provided a novel method to decipher the spatial domains by learning efficient representation from gene expressions and images via contrastive learning. The learned representations have been used efficiently for various downstream tasks, including trajectory inference, clustering, and visualization. This method will be particularly useful with the rapidly increasing spatial transcriptomics datasets.

### Key Points

- Existing methods for deciphering spatial domains only use the histopathological images to construct spot relations without updating in the training stage, or simply concatenate the information from gene expression and image into one feature vector.
- Here, we propose a novel method ConGI to accurately decipher spatial domains by integrating gene expression and histopathological images, where the gene expression is adapted to image information through contrastive learning. We introduce three contrastive loss functions within and between modalities to learn the common semantic representations across all modalities while avoiding their meaningless modality-private noise information.
- By comprehensive tests on tumor and normal spatial transcriptomics datasets, ConGI was shown to outperform existing methods in terms of spatial domain identification. More importantly, the representations learned from ConGI were used efficiently for various downstream tasks, including trajectory inference, clustering, and visualization.

## Code availability

All source codes used in our experiments have been deposited at https://github.com/biomed-AI/ConGI.

## Data availability

The spatial transcriptomics datasets that support the findings of this study are available here: (1) human HER2-positive breast tumor ST data https://github.com/almaan/HER2st/. (2) The LIBD human dorsolateral prefrontal cortex (DLPFC) data was acquired with 10X Visium composed of spatial transcriptomics data acquired from twelve tissue slices (http://research.libd.org/spatialLIBD/). (3) The mouse brain anterior section from 10X Visium (https://www.10xgenomics.com/resources/datasets). (4) the hum an epidermal growth factor receptor (HER) 2-amplified (HER+) invasive ductal carcinoma (IDC) (https://support.10xgenomics.com/spatial-gene-expression/datasets).

